# Genetic risk for coronary heart disease alters the influence of Alzheimer’s genetic risk on mild cognitive impairment

**DOI:** 10.1101/432443

**Authors:** Jeremy A. Elman, Matthew S. Panizzon, Mark W. Logue, Nathan A. Gillespie, Michael C. Neale, Chandra A. Reynolds, Daniel E. Gustavson, Ole A. Andreassen, Anders M. Dale, Carol E. Franz, Michael J. Lyons, William S. Kremen

## Abstract

**BACKGROUND:** Alzheimer’s disease (AD) is under considerable genetic influence. However, known susceptibility loci only explain a modest proportion of variance in disease outcomes. This small proportion could occur if the etiology of AD is heterogeneous. We previously found that an AD polygenic risk score (PRS) was significantly associated with mild cognitive impairment (MCI), an early stage of AD. Poor cardiovascular health is also associated with increased risk for AD and has been found to interact with AD pathology. Conditions such as coronary artery disease (CAD) are also heritable, and may contribute to heterogeneity if there are interactions of genetic risk for these conditions as there is phenotypically. However, case-control designs based on prevalent cases of a disease with relatively high case-fatality rate such as CAD may be biased toward individuals who have long post-event survival times and may therefore also identify loci with protective effects.

**METHODS:** We compared interactions between an AD-PRS and two CAD-PRSs, one based on a GWAS of incident cases and one on prevalent cases, on MCI status in 1,209 individuals.

**RESULTS:** As expected, the incidence-based CAD-PRS interacts with the AD-PRS to further increase MCI risk. Conversely, higher prevalence-based CAD-PRSs reduced the effect of AD genetic risk on MCI status.

**CONCLUSIONS:** These results demonstrate: i) the utility of including multiple PRSs and their interaction effects; ii) how genetic risk for one disease may modify the impact of genetic risk for another; and iii) the importance of considering ascertainment procedures of GWAS being used for genetic risk prediction.

## INTRODUCTION

Alzheimer’s disease (AD) is highly heritable(1), with the *APOE-ε4* allele having by far the greatest impact of any genetic locus. Large-scale genome-wide association studies (GWAS) of AD have identified 19 additional susceptibility loci(2), yet common variants identified by GWAS tend to account for only a small proportion of the variance in most complex diseases(3). The variance explained in AD risk can be increased using polygenetic risk score (PRS) approaches, which sum across many variants with small effect sizes(4). Our group further found that an AD-PRS is also associated with significantly higher odds of mild cognitive impairment (MCI)(5). These results lend support to the idea that MCI represents an early stage of AD, and demonstrate the utility of PRS in early identification. A study using a multiple polygenic risk score approach (including PRSs associated with multiple traits in a model) increased the proportion of explained variance in complex traits such as general cognitive ability(6), but this analysis did not examine the potential interactive effects of genetic risk factors or examine AD or MCI as an outcome. Rather than simply increasing the overall risk burden directly, it may be that certain additional genetic risk factors exert their effect by conferring additional susceptibility or resilience to the effects of primary AD risk genes.

Poorer cardiovascular health has been shown to be a significant risk factor for cognitive decline and progression to dementia(7–10), and vascular dementia is a common source of non-AD cognitive impairment. However, many patients demonstrate both AD and vascular lesions, and the presence of both greatly increases the odds of dementia(11, 12). Although some findings suggest that vascular and coronary risk are independent of Aβ pathology(13–15), others have found direct effects(16, 17). Whether amyloidogenic or not, vascular risk factors do appear to moderate the deleterious effects of AD pathology on cognitive and brain outcomes(18–20).

Coronary artery disease (CAD) is also under considerable genetic influence(21). Previous studies have found that the *APOE* and lipoprotein lipase genes are risk factors for both AD and CAD(22–24), suggesting some common biological basis. Genetic risk also appears to moderate the link between these diseases. For example, vascular risk factors increase the odds of cognitive decline or conversion to AD much more strongly in carriers of the *APOE-ε4* allele(25, 26). However, the extent to which additional susceptibility loci identified by GWAS interact is less clear. AD is a complex, polygenic disease. Thus, a model that incorporates PRSs for AD and CAD presents an opportunity to better characterize the potentially heterogenous genetic etiology of disease outcomes. Findings of synergistic effects at the phenotypic level between AD pathology and vascular risk further underscore the need to examine interactions of genetic risk for these factors in the context of multiple PRS models.

When generating a PRS, it is important to consider how the corresponding trait or disease status is defined in the original GWAS. The most common design for GWAS is case-control, which often depends on identifying prevalent cases. When the trait in question has a relatively high case-fatality rate, this may induce incidence-prevalence bias, also known as Neyman’s bias(27, 28). A GWAS of prevalent cases may be biased toward including individuals with lower mortality rates because individuals with shorter survival times after disease onset are less likely to be available for inclusion. Therefore, putative risk loci may actually be associated with increased survival time after disease onset in addition to those associated with disease onset itself. Incident cases of CAD would include individuals with both brief and extended post-event survival times(29), decreasing such bias. Thus, the loci detected in incidence-based versus prevalence-based analyses may represent somewhat different genetic influences(29), and may differently affect risk for AD or MCI.

In the present study, we examined how genetic risks for AD and CAD associate with MCI status in late middle-aged men. Better characterizing the genetic influences on this early disease stage may improve our ability to identify those individuals most appropriate for intervention. Based on evidence of phenotypic interactions between AD pathology and CAD risk factors, we focused on the interaction of genetic risk for AD and CAD. Importantly, to determine if the way in which cases were identified alters the association, we assessed one PRS based on prevalent cases of CAD and a second based on incident cases of CAD. Given that case-control designs of incident cases are less biased towards individuals with longer survival times, we predicted that an incident-based CAD-PRS would more strongly exacerbate the effect of AD genetic risk on cognitive status.

## METHODS & MATERIALS

### Participants

There were 1,329 men in the Vietnam Era Twin Study of Aging (VETSA)(30, 31) who were determined to be of white, non-Hispanic European ancestry (WNH). As PRSs are primarily ancestry specific, and large scale GWASs have been performed in WNH subjects, we excluded subjects of other ancestry from the analysis. We then excluded those with missing data that would preclude a possible MCI diagnosis, and with conditions that could cause cognitive deficits unrelated to MCI including seizure disorder, multiple sclerosis, stroke, HIV/AIDS, schizophrenia, substance dependence, or brain cancer(32). Additionally, in the present study the MCI group was limited to participants with amnestic MCI (aMCI). The final sample comprised 1,208 participants.

Sample characteristics are shown in **Table 1**. VETSA constitutes a national sample comparable to American men in their age range with respect to health and lifestyle characteristics(33). All were in some branch of military service sometime between 1965 and 1975. Nearly 80% report no combat exposure. VETSA participants had to be 51-59 years old at the time of recruitment in wave 1, and both twins in a pair had to be willing to participate(30, 31). Here we included wave 1 and new wave 2 participants, so that all were undergoing their initial assessment. In sum, VETSA constitutes a reasonably representative sample of community-dwelling men in their age range who were not selected for any health or diagnostic characteristic.

**Table 1.** Sample characteristics.

| Group | Cognitively Normal | Amnesic MCI |
| --- | --- | --- |
| N | 1119 | 89 |
| Age, *mean (SD)* | 56.7 (3.3) | 57.2 (3.5) |
| *APOE*-ε4+ | 29.4% | 26.2% |
| Ischemic Heart Disease* | 13.3% | 3.5% |
| Depressive symptoms, *mean (SD)* | 7.8 (7.6) | 9.0 (8.4) |
| Diabetes | 10.7% | 11.5% |

\*Ischemic heart disease variable is a summary measure that includes history of myocardial infarction, cardiac procedure or angina.

### Health/medical measures

A comprehensive medical history was collected for all participants(34). A summary measure of ischemic heart disease was created based on diagnosis or self-report of myocardial infarction, cardiac procedure (e.g. stent, balloon angioplasty, coronary artery bypass) or angina(35). Depressive symptoms were assessed with the Center for Epidemiological Studies Depression Scale(36). Diabetes was assessed if a participant reported being told by a physician that he had diabetes or if he was taking medication for diabetes. Type 1 diabetes would have ruled out entry into the military.

### Definition of mild cognitive impairment

MCI was diagnosed using the Jak-Bondi actuarial/neuropsychological approach(37, 38). Participants completed a comprehensive neuropsychological test battery comprising 18 tests covering 6 cognitive domains, as described elsewhere(32). To account for change from “premorbid” levels, we adjusted neuropsychological scores for a measure of young adult general cognitive ability(39, 40). Impairment in a cognitive domain was defined as having at least two tests that were >1.5 SDs below age- and education-adjusted normative means. The MCI group was restricted to individuals classified as amnestic MCI (aMCI; e.g., impaired memory domain). With this criterion, 1,119 (92.6%) individuals were cognitively normal (CN), and 89 (7.4%) individuals had aMCI. Individuals with non-amnestic MCI were not included in the analysis. Support for the validity of these criteria comes from our finding that higher AD-PRSs were associated with significantly increased odds of aMCI in these individuals(5).

### Genotyping methods

Genotyping and SNP cleaning methods have been described previously in detail(5), but are summarized here in brief. Whole genome genetic variation was assessed at deCODE Genetics (Reykjavík, Iceland). Genotyping was performed on Illumina HumanOmniExpress-24 v1.0A (Illumina, San Diego, CA). Beadchips were imaged using the Illumina iScan System and analyzed with Illumina GenomeStudio v2011.1 software containing Genotyping v1.9.4 module.

Cleaning and quality control of genome-wide genotype data was performed using PLINK v1.9(41). SNPs with more than 5% missing data or SNPs with Hardy-Weinberg equilibrium P-values <10^−6^ were excluded. Self-reported ancestry was confirmed using both SNPweights(42) and a principal components (PCs) analysis performed in PLINK v1.9 in conjunction with 1000 Genomes Phase 3 reference data(43). Analyses were restricted to participants of primarily European ancestry. PCs for use as covariates to control for population substructure were recomputed among this WNH set. Imputation was performed using MiniMac(44, 45) computed at the Michigan Imputation Server (https://imputationserver.sph.umich.edu). The 1,000 genomes phase 3 EUR data were used as a haplotype reference panel. Due to concerns about potential distortion in the haplotype-phasing step of imputation, only one randomly chosen participant’s data per genotyped MZ twin pair was submitted for imputation, and that participant’s resulting imputed data were applied to his MZ co-twin.

### Polygenic risk score calculation

The AD polygenic risk scores (AD-PRSs) were computed using summary data from the AD GWAS as presented in Lambert et al.(46). Individual SNP effect estimates and P-values were downloaded from http://web.pasteur-lille.fr/en/recherche/u744/igap/igap_download.php. Summary statistics from the coronary artery disease GWAS(47) used for the prevalent CAD-PRS have been contributed by CARDIoGRAMplusC4D investigators and have been downloaded from http://www.CARDIOGRAMPLUSC4D.ORG. The incident CAD-PRSs were computed using data from a GWAS on incident coronary heart disease(29) downloaded from the dbGaP web site, under phs000930.v6.p1 (https://www.ncbi.nlm.nih.gov/projects/gap/cgi-bin/study.cgi?study_id=phs000930.v6.p1).

Each PRS is a weighted average of VETSA sample additive imputed SNP dosages with the log-odds ratios (ORs) for each SNP estimated in the GWAS used as the weights. Rare SNPs (MAF<1%) and SNPs with poor imputation quality (R^2^<0.5) were excluded from PRS calculation. The remaining SNPs were trimmed for LD using PLINK’s clumping procedure (r^2^ threshold of 0.2 in a 500 kb window) based on LD patterns in the 1000 Genomes EUR cohort. PRSs were computed by PLINK v1.9 using a *P*-value threshold of *P*<0.50 for the AD-PRS because that threshold best differentiated AD or MCI cases from cognitively normal adults in 3 studies, including our own(4, 5, 48). The prevalence-based and incidence-based CAD-PRSs were both calculated with a threshold of *P*<0.05 because they showed the strongest association with the heart disease phenotype [incident CAD-PRS: *t*=2.631, *p*=0.001; prevalent CAD-PRS: *t*=3.690; *p*<0.001]. Genetic correlations between the 3 PRSs (AD, prevalent CAD, and incident CAD) were tested with LD Score regression software(49, 50) using the base summary statistics as input.

To determine whether interactions with the AD-PRS were being driven by the *APOE* locus or were independent of *APOE*, a second version of the AD-PRS was computed that excluded the region of LD surrounding the *APOE* gene (44,409,039 to 46,412,650 bp according to GRch37/Feb 2009). In models using this version of the AD-PRS, we additionally examined the influence of *APOE-ε4* measured by direct genotyping(51) separately from the AD-PRS.

### Statistical analysis

Differences in demographic variables were examined with chi-square tests and t-tests. We performed mixed effects logistic regression analyses using the glmer function from the *lme4* package(52) in R v3.2.1(53) to examine interactions between the AD-PRS and each CAD-PRS (i.e., incidence- and prevalence-based) on aMCI status. Although differentiating effects of *APOE* from other genes that contribute to the AD-PRS was not a primary focus of this study, we conducted secondary analyses to determine whether interaction effects were driven by the *APOE* gene. These analyses included two interactions: 1) the interaction between a given CAD-PRS and *APOE-ε4* carrier status, and 2) the interaction between a given CAD-PRS and the AD-PRS excluding the *APOE* region. All analyses adjusted for the first 3 PCs in order to account for any cryptic population substructure(54–56). We also adjusted for the following factors that may affect cognitive function: age, diabetes, and depressive symptoms (from the CESD), and history of head injury. Pair ID was included as a random effect to account for the non-independence within twin pairs.

## RESULTS

CN and aMCI groups did not differ with respect to age, *APOE-ε4* status, depressive symptoms, or diabetes (**Table 1**). There was a significantly greater proportion of individuals with ischemic heart disease in the CN group compared with the aMCI group [χ^2^ (1)=5.99, *p*=0.014]. There was a moderate genetic correlation between the incident CAD-PRS and prevalent CAD-PRS [r_g_=0.55; *p*=0.01]. However, the AD-PRS was not genetically correlated with either CAD-PRS [incident CAD-PRS: r_g_=0.04, *p*=0.89; prevalent CAD-PRS: r_g_=−0.06, *p*=0.55].

The model based on the AD-PRS and incident CAD-PRS showed main effects of both the AD-PRS [OR=1.54, *p*=0.003] and the incident CAD-PRS [OR=0.74, *p*=0.035]. There was also a significant *positive* interaction between the AD-PRS and the incident CAD-PRS [OR=1.42, *p*=0.009], with the association between the AD-PRS and aMCI status becoming stronger as incident CAD-PRSs increased. That is, as shown to the right of the dashed red line in **Figure 1A**, individuals at high genetic risk for AD were much more likely to have aMCI if they also had high genetic risk for incident CAD.

**Figure 1.**
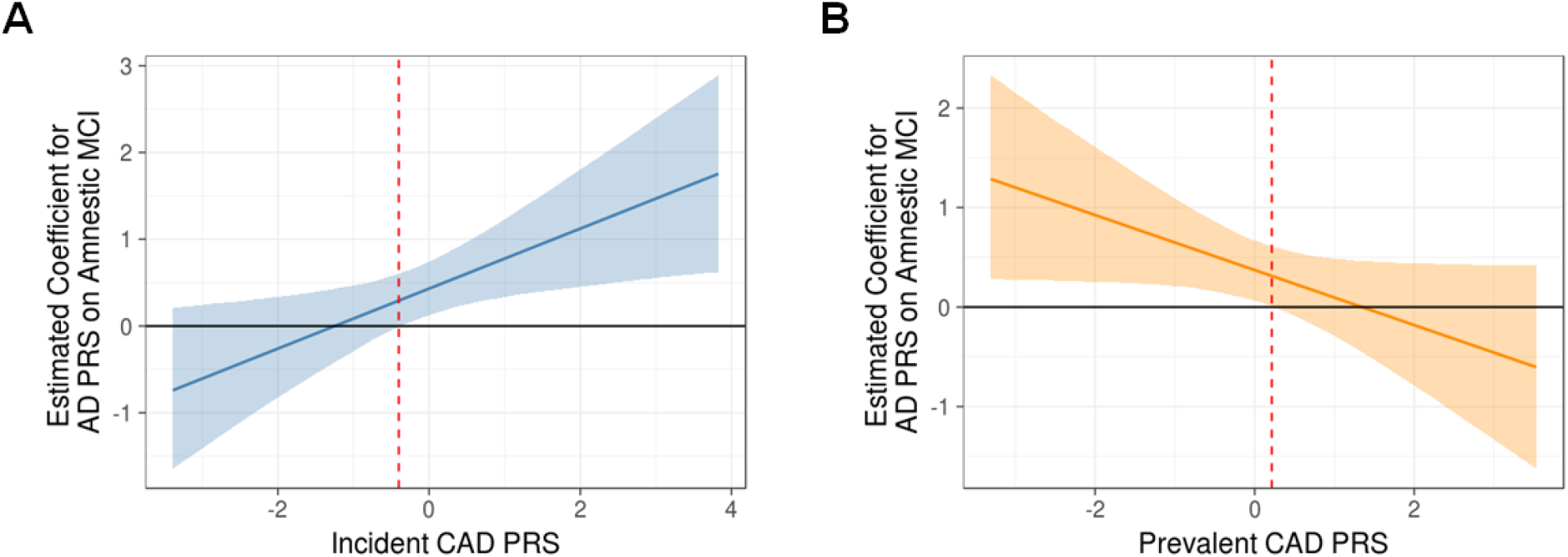
Interaction effects of polygenic risk scores for Alzheimer’s disease and coronary artery disease. Plots of the interaction of an Alzheimer’s disease polygenic risk score with A) a prevalent coronary artery disease polygenic risk score (CAD-PRS) and B) an incident CAD-PRS on amnestic mild cognitive impairment (MCI) status. The regression coefficient of the AD-PRS on amnestic MCI status is on the y-axis and is plotted across varying levels of CAD-PRSs on the x-axis. The dashed red line indicates the threshold of statistical significance for the AD-PRS as a predictor of aMCI status (i.e., where the 95% confidence intervals do not include 0). In 1A the AD-PRS is more predictive of risk for aMCI to the right of the dashed line (i.e., people with higher AD-PRSs are more likely to have aMCI if they also have *higher incident* CAD--RSs). In 1B the AD-PRS is a significant predictor of increased risk for aMCI to the left of the dashed line but is not significant to the right of the dashed line (i.e., people with higher AD-PRSs are only are higher risk for aMCI if they also have *lower prevalent* CAD-PRSs).

There was a very different result in the model based on the AD-PRS and the prevalent CAD-PRS. There was a significant main effect of the AD-PRS [OR=1.45, *p*=0.08] such that individuals with a higher score had greater odds of being in the aMCI group. There was no main effect of the prevalent CAD-PRS. However, there was a significant *negative* interaction between the AD-PRS and the prevalent CAD-PRS [OR=0.76, *p*=0.044], with the association between the AD-PRS and aMCI status weakening as prevalent CAD-PRSs increased. In other words, as shown to the left of the dashed red line in **Figure 1B**, the AD-PRS was significantly predictive of aMCI status when prevalent CAD-PRS scores were low, but no longer predictive when prevalent CAD-PRS scored were high.

We additionally tested models including separate interactions of the CAD-PRSs with both *APOE-ε4* status and the AD-PRS with *APOE* regions excluded. As before, in the model based on the incident CAD-PRS, both the main effect of the AD-PRS [OR=1.49, *p*=0.007] and the incident CAD-PRS [OR=0.69, *p*=0.021] remained significant. The interaction between the AD-PRS and the incident CAD-PRS [OR=1.31, p=0.048] remained significant as well. The AD-PRS was more strongly associated with increased risk of aMCI when the incident CAD-PRS was also high. The interaction between the incident CAD-PRS and *APOE* was not significant [OR=1.06, *p*=0.863].

The model based on the prevalent CAD-PRS showed a significant main effect of the AD-PRS [OR=1.39, *p*=0.020]. However, the interaction between the prevalent CAD-PRS and AD-PRS was reduced to a trend [OR=0.78, *p*=0.080] when the *APOE* region was excluded, indicating that the *APOE* gene was at least partially responsible for the interaction effect with the prevalent CAD-PRS.

## DISCUSSION

Here, we chose to examine PRSs for CAD in addition to an AD-PRS because CAD is an important risk factor for AD(7–10). More importantly, we examined whether there were interactive effects of genetic risk that mirror findings at the phenotypic level(18–20). Another report also included multiple PRSs to explain variance in complex traits(6), but that study differs from the present one in two key ways: 1) its PRSs were selected based on heritability rather than relationship to the outcome of interest; and 2) interactions between PRSs were not examined. We found that PRSs for CAD – a risk factor for AD – significantly moderated the association between genetic risk for AD and MCI status. Moreover, the interaction of the AD-PRS with the CAD-PRS went in opposite directions depending on whether the CAD-PRS was based on incident or prevalent cases. The association between the AD-PRS and an incidence-based CAD-PRS was positive, such that individuals at genetic risk for AD (i.e., high AD-PRS) were even more likely to have MCI when they also had a high incident CAD-PRS. In contrast, there was a somewhat counterintuitive interaction between the AD-PRS and a prevalence-based CAD-PRS. This interaction was negative, such that the AD-PRS was predictive of MCI when scores on the prevalent CAD-PRS were low, but no longer predictive of MCI when score on the CAD-PRS were high.

These results illustrate the usefulness of testing interactions between PRSs on complex traits. The genetic underpinnings of AD are multifactorial, with significant risk loci linked to various biological pathways(57, 58). Thus, individuals may progress to AD along multiple routes and this progression may be further mitigated or exacerbated by various other factors. Incorporating multiple risk factor PRSs and their interactions may capture the genetic etiology of AD more fully and help explain variability in the relationship between genetic risk for AD and clinical outcomes. When examining only main effects in the current study, it would appear that genetic risk for CAD was either not associated (prevalent CAD-PRS), or even negatively associated (incident CAD-PRS) with risk of MCI. Yet the significant interactions illustrate how additional genetic factors may exert their influence by moderating the relationship between primary AD risk genes and disease outcomes.

Genetic loci identified in GWAS of both incident and prevalent cases of CAD should be associated with poor cardiovascular health. Potential mechanisms for this added risk are that vascular factors such as hypertension can weaken the blood brain barrier, exposing the brain to harmful systemic elements(10); vascular risk factors may contribute to formation or disrupt clearance of amyloid(59, 60); and vascular risk factors may potentiate the toxic effects of amyloid on brain tissue(19). Individuals with a high incident CAD-PRS may therefore have cardiovascular systems more vulnerable to AD-related pathological processes.

The seemingly protective effects of the prevalence-based CAD-PRS and the higher rate of ischemic heart disease among cognitively normal participants compared to MCI may seem counterintuitive. However, a potential explanation for this is the incidence-prevalence (or Neyman) bias(27, 29). When including prevalent cases in a case-control design of a disease with relatively high case-fatality rates, the sample will be inherently biased toward individuals that survive. Individuals with CAD that lived long enough to be identified for a GWAS of prevalent cases may be more resilient to cardiovascular insult, with some of this resilience arising from genetic factors. Likewise, individuals with ischemic heart disease in the VETSA sample were the subset of cases that not only survived a cardiac event, but were healthy enough to travel and participate in the study. It has been proposed that some of the neurodegeneration and associated cognitive decline in AD may be caused by disruptions to cerebral microvasculature, and that this damage can mirror changes to systemic vasculature(61, 62). Therefore, genetic influences conferring resilience against the effects of cardiovascular events may also protect against cognitive decline and would explain the negative interaction found here. A similar argument has been made to explain why smoking can be negatively associated with prevalent cases of AD (i.e., an apparent protective effect), but positively associated with incident cases of AD (indicating it is a risk factor)(63). The results of the present study should therefore not be taken to suggest that the onset of CHD is a protective factor against cognitive decline. Rather, those individuals who have long survival times following a cardiovascular event may be more resilient to both vascular damage and cognitive decline due to genetic or other protective factors. It is the genetic influences that confer resilience in the face of cardiovascular events—not genetic influences on cardiovascular disease itself—that have some protective effects. Looked at from the other direction, the negative interaction indicated that the AD-PRS was predictive of MCI for people with low prevalence-based CAD-PRSs. This should not be taken to mean that low CAD risk increases risk for MCI or AD. Rather, it suggests that in the absence of other risk factors, AD risk alleles alone play a greater role in risk for developing MCI or AD.

The primary focus of the present study was not to dissociate effects of *APOE* from other AD risk loci, but there were nevertheless some interesting findings. The interaction of the incident CAD-PRS was not specific to *APOE*, whereas the negative interaction of the prevalent CAD-PRS with genetic risk for AD appeared to be weakened when the *APOE* genotype was included separately. When separated out, the interaction with the AD-PRS (excluding the *APOE* region) was no longer significant. This is consistent with previous findings that the *APOE* gene and the genes comprising the AD-PRS may be differentially associated with different traits such as amyloid deposition, hippocampal volume, and cognition(64). It is perhaps not surprising that there would be some links between a CAD-PRS and *APOE* given that the *APOE-ε4* allele is itself a risk factor for CAD, and that vascular risk factors are more strongly related to cognitive decline among APOE-ε4 carriers(22, 23, 25, 26).

Interestingly, death from CAD appears to be heritable(65) and at least some of this risk may be attributable to the *APOE* gene. *APOE* has been proposed as a “frailty gene”, with the ε4 allele associated with increased mortality risk at younger ages(66), and specifically with higher mortality in cases of CAD(67, 68). This effect on mortality is strongest during middle age, the age at which VETSA participants were assessed in this study, and weakens at older ages(69). The incidence-prevalence bias may therefore be exacerbated in individuals at genetic risk for both AD and CAD. That is, individuals with high genetic risk for both diseases may be even less likely to survive long enough to be captured in case-control designs of prevalent CAD after cardiac events, contributing to an apparent negative interaction between these two genetic risk factors.

There are a few limitations to this study. The first is that the VETSA is an all-male sample, and therefore these particular results may not generalize to women. However, there is no reason to believe that the potential for interactions of genetic risk occur in men but not women. Second, our MCI diagnosis was not confirmed using biomarkers. However, the Jak-Bondi actuarial/neuropsychological approach has been well-validated and performs favorably compared to other MCI classification schemes with regards to biomarker positivity, clinical progression, and reduced rates of reversion to cognitively normal. Moreover, we previously showed individuals diagnosed as MCI with this approach have higher genetic risk for AD (also indicated by the significant main of the AD PRS in the current analysis). It is also important to note that the presence or absence of biomarker confirmation does not alter the interpretation of the key finding here that genetic risk for one disease may modify the impact of genetic risk for another.

The current study raises three important points. The first is that examining interactive effects of multiple PRSs can further explain variability in the association between genetic risk for AD and cognitive outcomes, even when main effects may be absent. Complex traits such as AD are likely to have a heterogenous genetic basis and the impact of primary risk loci may be moderated by separate genetic factors. Thus, more fully describing this variability will aid in identifying individuals most at risk and help predict the likelihood and/or rate of disease progression. Second, while it is important to examine interactions with the *APOE* risk locus because the *APOE-ε4* allele is the largest single genetic determinant of AD risk, a greater focus on interaction effects between PRSs is warranted given the polygenic nature of AD. Third, the design of the base GWAS used to calculate PRSs must be considered to appropriately interpret what traits the effect alleles actually represent, particularly when there is a high case-fatality rate. As shown here, this can even result in the reversal of expected effects, with susceptibility loci demonstrating a protective moderating effect on genetic risk for a given disease. Future work incorporating longitudinal follow-ups will be necessary to determine whether individuals with varying degrees of genetic risk for AD and its related risk factors demonstrate clearly dissociable patterns of disease progression.

## ACKNOWLEDGMENTS

This work was supported by National Institute on Aging R01 AG050595 (W.S.K., M.J.L., C.E.F.), R01 AG022381 (W.S.K.), R03 AG046413 (C.E.F), and K08 AG047903 (M.S.P), Research Council of Norway (223273) (O.A.A.), Stiftelsen KG Jebsen (O.A.A.), and the VA San Diego Center of Excellence for Stress and Mental Health Healthcare System. The content is the responsibility of the authors and does not necessarily represent official views of the NIA, NIH, or VA. The Cooperative Studies Program of the U.S. Department of Veterans Affairs provided financial support for development and maintenance of the Vietnam Era Twin Registry. We would also like to acknowledge the continued cooperation and participation of the members of the VET Registry and their families.

## REFERENCES

1. Gatz M, Reynolds CA, Fratiglioni L, Johansson B, Mortimer JA, Berg S, et al. (2006): Role of genes and environments for explaining Alzheimer disease. Arch Gen Psychiatry. 63:168–174.

2. Lambert JC, Ibrahim-Verbaas CA, Harold D, Naj AC, Sims R, Bellenguez C, et al. (2013): Meta-analysis of 74,046 individuals identifies 11 new susceptibility loci for Alzheimer’s disease. Nat Genet. 45:1452–1458.

3. Visscher PM, Brown MA, McCarthy MI, Yang J (2012): Five years of GWAS discovery. Am J Hum Genet. 90:7–24.

4. Escott-Price V, Sims R, Bannister C, Harold D, Vronskaya M, Majounie E, et al. (2015): Common polygenic variation enhances risk prediction for Alzheimer’s disease. Brain. 138:3673–3684.

5. Logue MW, Panizzon MS, Elman JA, Gillespie NA, Hatton SN, Gustavson DE, et al. (2018): Use of an Alzheimer’s disease polygenic risk score to identify mild cognitive impairment in adults in their 50s. Mol Psychiatry.

6. Krapohl E, Patel H, Newhouse S, Curtis CJ, von Stumm S, Dale PS, et al. (2018): Multi-polygenic score approach to trait prediction. Mol Psychiatry. 23:1368–1374.

7. Jefferson AL, Beiser AS, Himali JJ, Seshadri S, O’Donnell CJ, Manning WJ, et al. (2015): Low cardiac index is associated with incident dementia and Alzheimer disease: the Framingham Heart Study. Circulation. 131:1333–1339.

8. Viticchi G, Falsetti L, Buratti L, Boria C, Luzzi S, Bartolini M, et al. (2015): Framingham risk score can predict cognitive decline progression in Alzheimer’s disease. Neurobiology of Aging. 36:2940–2945.

9. Kaffashian S, Dugravot A, Elbaz A, Shipley MJ, Sabia S, Kivimaki M, et al. (2013): Predicting cognitive decline: a dementia risk score vs. the Framingham vascular risk scores. Neurology. 80:1300–1306.

10. Skoog I (2000): Vascular aspects in Alzheimer’s disease. Journal of neural transmission Supplementum. 59:37–43.

11. Azarpazhooh MR, Avan A, Cipriano LE, Munoz DG, Sposato LA, Hachinski V (2018): Concomitant vascular and neurodegenerative pathologies double the risk of dementia. Alzheimers Dement. 14:148–156.

12. Jellinger KA (2008): Morphologic diagnosis of “vascular dementia” - a critical update. J Neurol Sci. 270:1–12.

13. Esiri MM, Joachim C, Sloan C, Christie S, Agacinski G, Bridges LR, et al. (2014): Cerebral subcortical small vessel disease in subjects with pathologically confirmed Alzheimer disease: a clinicopathologic study in the Oxford Project to Investigate Memory and Ageing (OPTIMA). Alzheimer disease and associated disorders. 28:30–35.

14. Marchant NL, Reed BR, Sanossian N, Madison CM, Kriger S, Dhada R, et al. (2013): The aging brain and cognition: contribution of vascular injury and abeta to mild cognitive dysfunction. JAMA Neurol. 70:488–495.

15. Vemuri P, Lesnick TG, Przybelski SA, Knopman DS, Preboske GM, Kantarci K, et al. (2015): Vascular and amyloid pathologies are independent predictors of cognitive decline in normal elderly. Brain. 138:761–771.

16. Reed BR, Marchant NL, Jagust WJ, DeCarli CC, Mack W, Chui HC (2012): Coronary risk correlates with cerebral amyloid deposition. Neurobiol Aging. 33:1979–1987.

17. van Norden AG, van Dijk EJ, de Laat KF, Scheltens P, Olderikkert MG, de Leeuw FE (2012): Dementia: Alzheimer pathology and vascular factors: from mutually exclusive to interaction. Biochim Biophys Acta. 1822:340–349.

18. Brickman AM, Honig LS, Scarmeas N, Tatarina O, Sanders L, Albert MS, et al. (2008): Measuring cerebral atrophy and white matter hyperintensity burden to predict the rate of cognitive decline in Alzheimer disease. Arch Neurol. 65:1202–1208.

19. Villeneuve S, Reed BR, Madison CM, Wirth M, Marchant NL, Kriger S, et al. (2014): Vascular risk and Aβ interact to reduce cortical thickness in AD vulnerable brain regions. Neurology. 83:40–47.

20. Snowdon DA, Greiner LH, Mortimer JA, Riley KP, Greiner PA, Markesbery WR (1997): Brain infarction and the clinical expression of Alzheimer disease. The Nun Study. JAMA. 277:813–817.

21. Zdravkovic S, Wienke A, Pedersen NL, Marenberg ME, Yashin AI, De Faire U (2002): Heritability of death from coronary heart disease: a 36-year follow-up of 20 966 Swedish twins. J Intern Med. 252:247–254.

22. Eichner JE, Dunn ST, Perveen G, Thompson DM, Stewart KE, Stroehla BC (2002): Apolipoprotein E polymorphism and cardiovascular disease: a HuGE review. Am J Epidemiol. 155:487–495.

23. Song Y, Stampfer MJ, Liu S (2004): Meta-analysis: apolipoprotein E genotypes and risk for coronary heart disease. Ann Intern Med. 141:137–147.

24. Xie C, Wang ZC, Liu XF, Yang MS (2010): The common biological basis for common complex diseases: evidence from lipoprotein lipase gene. Eur J Hum Genet. 18:3–7.

25. Hofman A, Ott A, Breteler MM, Bots ML, Slooter AJ, van Harskamp F, et al. (1997): Atherosclerosis, apolipoprotein E, and prevalence of dementia and Alzheimer’s disease in the Rotterdam Study. Lancet. 349:151–154.

26. Helzner EP, Luchsinger JA, Scarmeas N, Cosentino S, Brickman AM, Glymour MM, et al. (2009): Contribution of vascular risk factors to the progression in Alzheimer disease. Arch Neurol. 66:343–348.

27. Hill G, Connelly J, Hebert R, Lindsay J, Millar W (2003): Neyman’s bias re-visited. J Clin Epidemiol. 56:293–296.

28. Neyman J (1955): Statistics - Servant of All Sciences. Science. 122:401–406.

29. Dehghan A, Bis Jc, White CC, Smith AV, Morrison AC, Cupples LA, et al. (2016): Genome-Wide Association Study for Incident Myocardial Infarction and Coronary Heart Disease in Prospective Cohort Studies: The CHARGE Consortium. PLoS One. 11:e0144997.

30. Kremen WS, Franz CE, Lyons MJ (2013): VETSA: The Vietnam Era Twin Study of Aging. Twin Res Hum Genet. 16:399–402.

31. Kremen WS, Thompson-Brenner H, Leung YJ, Grant MD, Franz CE, Eisen SA, et al. (2006): Genes, environment, and time: The Vietnam Era Twin Study of Aging (VETSA). Twin Res Hum Genet. 9:1009–1022.

32. Kremen WS, Jak AJ, Panizzon MS, Spoon KM, Franz CE, Thompson WK, et al. (2014): Early identification and heritability of mild cognitive impairment. Int J Epidemiol. 43:600–610.

33. Schoeneborn CA, Heyman KM (2009): Health characteristics of adults aged 55 years and over: United States, 2004-2007. National Health Statistics Reports. Hyattsville, MD: National Center for Health Statistics.

34. Kremen WS, Thompson-Brenner H, Leung YM, Grant MD, Franz CE, Eisen SA, et al. (2006): Genes, environment, and time: the Vietnam Era Twin Study of Aging (VETSA). Twin Res Hum Genet. 9:1009–1022.

35. Xian H, Scherrer JF, Franz CE, McCaffery J, Stein PK, Lyons MJ, et al. (2010): Genetic vulnerability and phenotypic expression of depression and risk for ischemic heart disease in the Vietnam era twin study of aging. Psychosom Med. 72:370–375.

36. Radloff LS (1977): The CES-D scale: A self-report depression scale for research in the general population. Applied Psychological Measurement. 1:385–401.

37. Bondi MW, Edmonds EC, Jak AJ, Clark LR, Delano-Wood L, McDonald CR, et al. (2014): Neuropsychological criteria for mild cognitive impairment improves diagnostic precision, biomarker associations, and progression rates. J Alzheimers Dis.

38. Jak AJ, Bondi MW, Delano-Wood L, Wierenga C, Corey-Bloom J, Salmon DP, et al. (2009): Quantification of five neuropsychological approaches to defining mild cognitive impairment. The American journal of geriatric psychiatry: official journal of the American Association for Geriatric Psychiatry. 17:368–375.

39. Lyons MJ, York TP, Franz CE, Grant MD, Eaves LJ, Jacobson KC, et al. (2009): Genes determine stability and the environment determines change in cognitive ability during 35 years of adulthood. Psychol Sci. 20:1146–1152.

40. Lyons MJ, Panizzon MS, Liu W, McKenzie R, Bluestone NJ, Grant MD, et al. (2017): A longitudinal twin study of general cognitive ability over four decades. Dev Psychol. 53:11701–177.

41. Chang CC, Chow CC, Tellier LC, Vattikuti S, Purcell SM, Lee JJ (2015): Second-generation PLINK: rising to the challenge of larger and richer datasets. GigaScience. 4:1–16.

42. Chen CY, Pollack S, Hunter DJ, Hirschhorn JN, Kraft P, Price AL (2013): Improved ancestry inference using weights from external reference panels. Bioinformatics. 29:1399–1406.

43. 1000 Genomes Project Consortium, Auton A, Brooks LD, Durbin RM, Garrison EP, Kang HM, et al. (2015): A global reference for human genetic variation. Nature. 526:68–74.

44. Howie B, Fuchsberger C, Stephens M, Marchini J, Abecasis GR (2012): Fast and accurate genotype imputation in genome-wide association studies through pre-phasing. Nat Genet. 44:955–959.

45. Fuchsberger C, Abecasis GR, Hinds DA (2015): minimac2: faster genotype imputation. Bioinformatics. 31:782–784.

46. Lambert JC, Ibrahim-Verbaas CA, Harold D, Naj AC, Sims R, Bellenguez C, et al. (2012): Meta-analysis of 74,046 individuals identifies 11 new susceptibility loci for Alzheimer’s disease. Nat Genet.

47. Nikpay M, Goel A, Won HH, Hall LM, Willenborg C, Kanoni S, et al. (2015): A comprehensive 1,000 Genomes-based genome-wide association meta-analysis of coronary artery disease. Nat Genet. 47:1121–1130.

48. Escott-Price V, Myers AJ, Huentelman M, Hardy J (2017): Polygenic Risk Score Analysis of Pathologically Confirmed Alzheimer’s Disease. Ann Neurol.n/a-n/a.

49. Bulik-Sullivan B, Finucane HK, Anttila V, Gusev A, Day FR, Loh PR, et al. (2015): An atlas of genetic correlations across human diseases and traits. Nat Genet. 47:1236–1241.

50. Bulik-Sullivan BK, Loh PR, Finucane HK, Ripke S, Yang J, Schizophrenia Working Group of the Psychiatric Genomics C, et al. (2015): LD Score regression distinguishes confounding from polygenicity in genome-wide association studies. Nat Genet. 47:291–295.

51. Schultz MR, Lyons mJ, Franz CE, Grant MD, Boake C, Jacobson KC, et al. (2008): Apolipoprotein E genotype and memory in the sixth decade of life. Neurology. 70:1771–1777.

52. Bates D, Maechler M, Bolker B, Walker S, Christensen RHB, Singmann H, et al. (2015): lme4: Linear Mixed-Effects Models using ‘Eigen’ and S4. 1.1-8 ed.

53. R Core Team (2017): R: A language and environment for statistical computing. Vienna, Austria: R Foundation for Statistical Computing.

54. Sadeh N, Spielberg JM, Logue MW, Wolf EJ, Smith AK, Lusk J, et al. (2016): SKA2 methylation predicts reduced cortical thickness in prefrontal cortex. Mol Psychiatry. 21:299.

55. Sadeh N, Wolf EJ, Logue MW, Lusk J, Hayes JP, McGlinchey RE, et al. (2016): Polygenic risk for externalizing psychopathology and executive dysfunction in trauma-exposed veterans. Clin Psychol Sci. 4:545–558.

56. Wolf EJ, Logue MW, Hayes JP, Sadeh N, Schichman SA, Stone A, et al. (2016): Accelerated DNA methylation age: Associations with PTSD and neural integrity. Psychoneuroendocrinology. 63:155–162.

57. International Genomics of Alzheimer’s Disease C (2015): Convergent genetic and expression data implicate immunity in Alzheimer’s disease. Alzheimers Dement. 11:658–671.

58. Sun Q, Xie N, Tang B, Li R, Shen Y (2017): Alzheimer’s Disease: From Genetic Variants to the Distinct Pathological Mechanisms. Front Mol Neurosci. 10:319.

59. Gottesman RF, Schneider AL, Zhou Y, Coresh J, Green E, Gupta N, et al. (2017): Association Between Midlife Vascular Risk Factors and Estimated Brain Amyloid Deposition. JAMA. 317:1443–1450.

60. Ramanathan A, Nelson AR, Sagare AP, Zlokovic BV (2015): Impaired vascular-mediated clearance of brain amyloid beta in Alzheimer’s disease: the role, regulation and restoration of LRP1. Front Aging Neurosci. 7:136.

61. Farkas E, De Vos RAI, Steur ENHJ, Luiten PGM (2000): Are Alzheimer’s disease, hypertension, and cerebrocapillary damage related? Neurobiology of Aging. 21:235–243.

62. Farkas E, Luiten PG (2001): Cerebral microvascular pathology in aging and Alzheimer’s disease. Prog Neurobiol. 64:575–611.

63. Wang H-X, Fratiglioni L, Frisoni GB, Viitanen M, Winblad B (1999): Smoking and the Occurence of Alzheimer’s Disease: Cross-Sectional and Longitudinal Data in a Population-based Study. American Journal of Epidemiology. 149:640–644.

64. Ge T, Sabuncu MR, Smoller JW, Sperling RA, Mormino EC, Alzheimer’s Disease Neuroimaging I (2018): Dissociable influences of APOE epsilon4 and polygenic risk of AD dementia on amyloid and cognition. Neurology. 90:e1605–e1612.

65. Marenberg ME, Risch N, Berkman LF, Floderus B, de Faire U (1994): Genetic susceptibility to death from coronary heart disease in a study of twins. The New England journal of medicine. 330:1041–1046.

66. Gerdes LU, Jeune B, Ranberg KA, Nybo H, Vaupel JW (2000): Estimation of apolipoprotein E genotype-specific relative mortality risks from the distribution of genotypes in centenarians and middle-aged men: apolipoprotein E gene is a “frailty gene,” not a “longevity gene”. Genet Epidemiol. 19:202–210.

67. Rosvall L, Rizzuto D, Wang HX, Winblad B, Graff C, Fratiglioni L (2009): APOE-related mortality: effect of dementia, cardiovascular disease and gender. Neurobiol Aging. 30:1545–1551.

68. Stengard JH, Weiss KM, Sing CF (1998): An ecological study of association between coronary heart disease mortality rates in men and the relative frequencies of common allelic variations in the gene coding for apolipoprotein E. Hum Genet. 103:234–241.

69. Ewbank DC (2002): Mortality differences by APOE genotype estimated from demographic synthesis. Genet Epidemiol. 22:146–155.

